# Minimalistic 3D Chromatin Models: Sparse Interactions in Single Cells Drive the Chromatin Fold and Form Many-Body Units

**DOI:** 10.1101/2021.08.06.455444

**Authors:** Jie Liang, Alan Perez-Rathke

## Abstract

Computational modeling of 3D chromatin plays an important role in understanding the principles of genome organization. We discuss methods for modeling 3D chromatin structures, with focus on a minimalistic polymer model which inverts population Hi-C into high-resolution, high-coverage single-cell chromatin conformations. Utilizing only basic physical properties such as nuclear volume and no adjustable parameters, this model uncovers a few specific Hi-C interactions (15-35 for enhancer-rich loci in human cells) that can fold chromatin into individual conformations consistent with single-cell imaging, Dip-C, and FISH-measured genomic distance distributions. Aggregating an ensemble of conformations also reproduces population Hi-C interaction frequencies. Furthermore, this single-cell modeling approach allows quantification of structural heterogeneity and discovery of specific many-body units of chromatin interactions. This minimalistic 3D chromatin polymer model has revealed a number of insights: 1) chromatin scaling rules are a result of volume-confined polymers; 2) TADs form as a byproduct of 3D chromatin folding driven by specific interactions; 3) chromatin folding at many loci is driven by a small number of specific interactions; 4) cell subpopulations equipped with different chromatin structural scaffolds are developmental stage-dependent; and 5) characterization of the functional landscape and epigenetic marks of many-body units which are simultaneously spatially co-interacting within enhancer-rich, euchromatic regions. The implications of these findings in understanding the genome structure-function relationship are also discussed.

## Introduction

Chromosome conformation capture (1–4) and imaging analysis (5–8) have generated a wealth of information on nuclear genome organization. Structural units, such as compartments, topologically associating domains (TADs), and loops, have been uncovered from block patterns in Hi-C frequency heatmaps (9–11). However, canonical Hi-C measures only population-averaged, pairwise interaction frequencies, which do not fully represent the underlying 3D conformational distributions in individual cells. This limits our understanding, as single-cell 3D chromatin structures do not automatically follow from Hi-C’s population-averaged heatmap representation (12).

Veiled behind each Hi-C heatmap is a population of 3D chromatin conformations. While single-cell technologies can provide direct information on 3D spatial arrangements of genomic elements within a nucleus (5, 11), they are limited in both resolution and coverage. As an alternative, computational modeling of chromatin polymers provides a powerful means for uncovering 3D spatial configurations of genomic regions and plays important roles in deciphering physical principles of genome organization.

### Optimization-Based 3D Chromatin Models

Among the many different approaches to modeling 3D chromatin from Hi-C, optimization methods aim to generate chromatin conformations maximally satisfying Hi-C derived distance-restraints (13–18) (reviewed in (19, 20)). However, they are *ad hoc* as conformations obtained do not follow a physically-governed, *a priori*-defined distribution, owing to the lack of a physical model underpinning these methods. In addition, there is no consideration to ensure adequate sampling such that a diverse structural ensemble is sufficiently represented. Furthermore, many tunable parameters are often employed to ensure a good fit to experimental data: as many distance thresholds as the number of restraints modeled may need to be adjusted. If inadequately regularized, they may lead to overfitting.

Understanding the spatial organization of chromatin requires a physical 3D chromatin model. With an accurate physical model, the sampled conformational ensemble will correctly reflect the conformational distribution in the cell population.

### 3D Heteropolymer Models with an Empirical Energy Function

We now briefly discuss physical models of heteropolymers with binding interactions among regions of different chromatin states (see excellent reviews of (21–23) for details). Here chromatin is modeled as heterogeneous blocks of 3D monomer chains. Monomers, representing 0.5 – 500 kb chromatin, are grouped into different blocks by their epigenetically-defined chromatin states. An empirical energy function describes how these monomers interact according to their binding affinities (24–31). Molecular dynamics simulations then generate an ensemble of chromatin conformations. While realistic dynamics are not possible at this coarsegrained scale, the ensembles obtained can provide direct 3D structural information.

These powerful models allow detailed assessment of the effects of different mechanistic assumptions, which can reveal important insights into principles of genome organization. As an example, studies based on the MiChroM method suggest the likely origin of chromosome territories, reproduce phase separation, and provide further evidence for the preferential localization of active genes (26, 27, 32). It was shown that epigenetic information can be used to predict the structural ensembles of multiple human cell lines, and short segments of chromatin make state transitions between closed conformations and open dumbbell conformations (33). The aggregation of denser and predominantly inactive chromatin was found to drive the positioning of active chromatin toward the surface of individual chromosomal territories (33). Recent efforts in scaling up MiChroM showed that inter-chromosomal interactions can now be studied through detailed molecular dynamics simulations (34). A study of the full diploid genome of all 46 human chromosomes at 1 Mb resolution was recently carried out to determine factors important for radial positioning of chromosomes (30). A separate study using a polymer model with only two epigenetic states and fixed loop anchors showed that compartments in Hi-C maps are due to microphase separation of these two states, leading to highly heterogeneous chromosome dynamics (28).

**Fig. 1.**
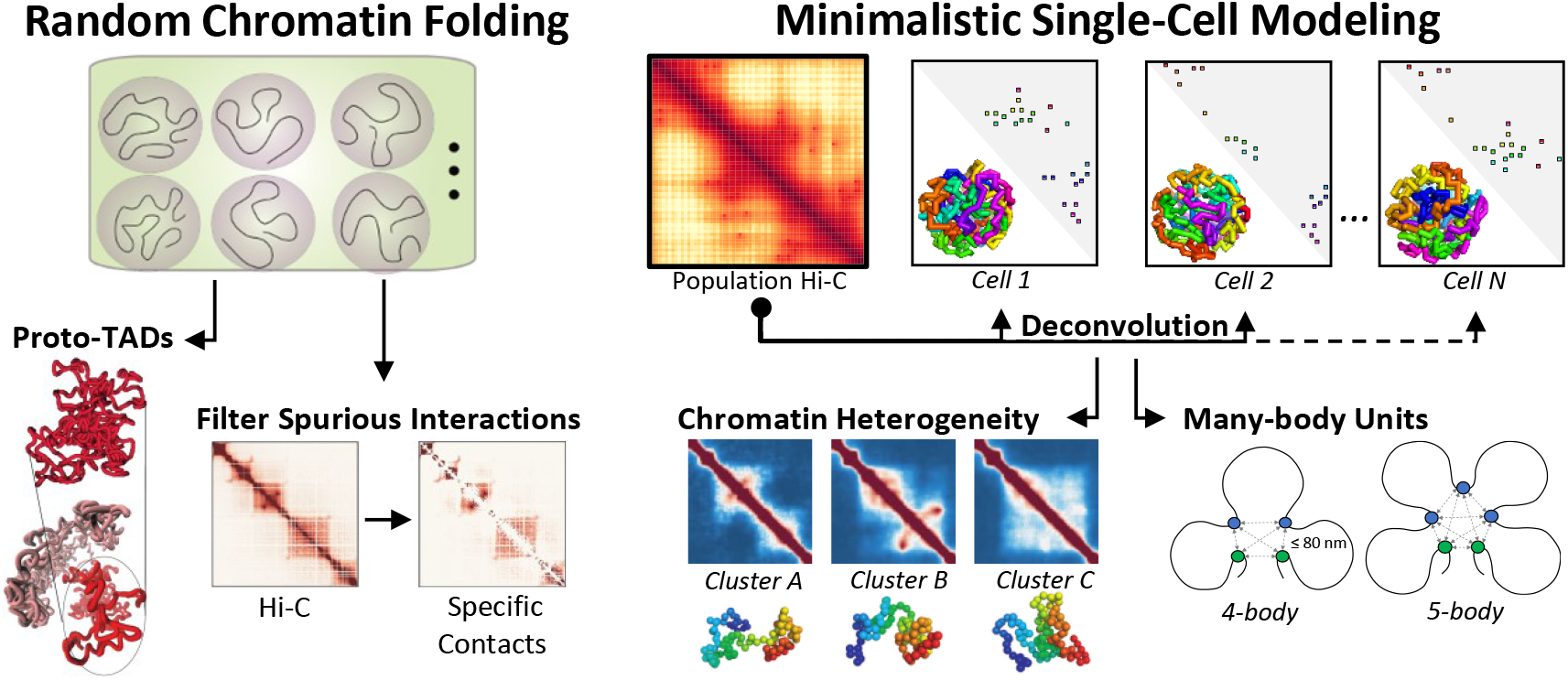
Graphic Abstract.

The String and Binders Switch (SBS) model and related methods (24) have been applied to study how binder mediated chromatin interactions can lead to 3D chromatin structures. In (35), the 3D structures of the *Sox9* locus and the whole mouse chromosome 7 were constructed. In (31), it was found that an increase in binder concentration can lead chromatin to adopt a coil-to-globule phase-separated state, where the intrinsically degenerate globule state corresponds to the large variation observed in modeled 3D chromatin conformations. Another recent study applied the SBS model to extract 3D chromatin structures from GAM, SPRITE and Hi-C data (36), and compared these experimental methods with *in silico* models. These experimental techniques were found to have different efficiency in capturing long-range interactions, requiring a different minimum number of cells (ranging from 250 to 800) for replicate experiments to return statistically consistent measurements (36).

### Binding Factors and Affinities

The behavior of heteropolymer chromatin is fully determined by the energy landscape of the model (24–28). Unlike molecular mechanics, where physical forces such as electrostatic interactions have been thoroughly studied, first-principle understanding of the physical factors and interactions of chromatin at mesoscale is not available. There are likely over 2,600 different proteins expressed in a cell (37), many yet to be identified, which complex with DNA and each other at often unknown rates and affinities. Instead, coarse-grained chromatin states and empirical binding affinities must be inferred from Hi-C and epigenetic data. There are likely many different chromatin state and binding affinity assignments that are all consistent with experimental data. The non-identifiable nature of such assignments, combined with other assumptions of the energy model, may hinder precise inference and limit the biological interpretability in phenomena detected through simulations. Furthermore, while an *a priori* constructed energy model can effectively explore consequences of various model assumptions, making biological discoveries not encoded in the model input is challenging.

### Importance of Thorough Sampling

A prerequisite for all 3D chromatin polymer methods is that conformational ensembles must be thoroughly sampled. This is exceedingly challenging as chromosomes are severely confined in the nuclear volume (38) and exhibit extraordinary heterogeneity (5, 6, 33, 39). Generating biologically-accurate chromatin ensembles using molecular dynamics is non-trivial. Without thorough sampling, it is difficult to ascertain if bias is present due to inadequate sampling, a misspecified energy model, or both.

## Minimalistic Self-Avoiding Polymer Model of 3D Chromatin

Another approach is to model chromatin as a 3D self-avoiding polymer but with minimal physical properties and no adjustable parameters. Once its emerging behavior is characterized and deficiencies identified, additional ingredi-ents are then introduced to refine the model. Such a minimalistic approach of polymer modeling has had great success in earlier studies of protein folding (40, 41).

The initial premises are that chromatin must be 1) connected, 2) self-avoiding, and 3) confined in the cell nucleus. This approach becomes feasible with recent deep-sampling algorithms (38, 42–45). These algorithms generate nuclear-confined, self-avoiding chromatin polymers by sequentially placing connected monomer units until the target polymer length is reached (Fig 2a). Advanced sampling techniques such as fractal Monte Carlo enable generation of large and complex ensembles consisting of 10^4-5^ single-cell 3D chromatin structures.

**Fig. 2.**
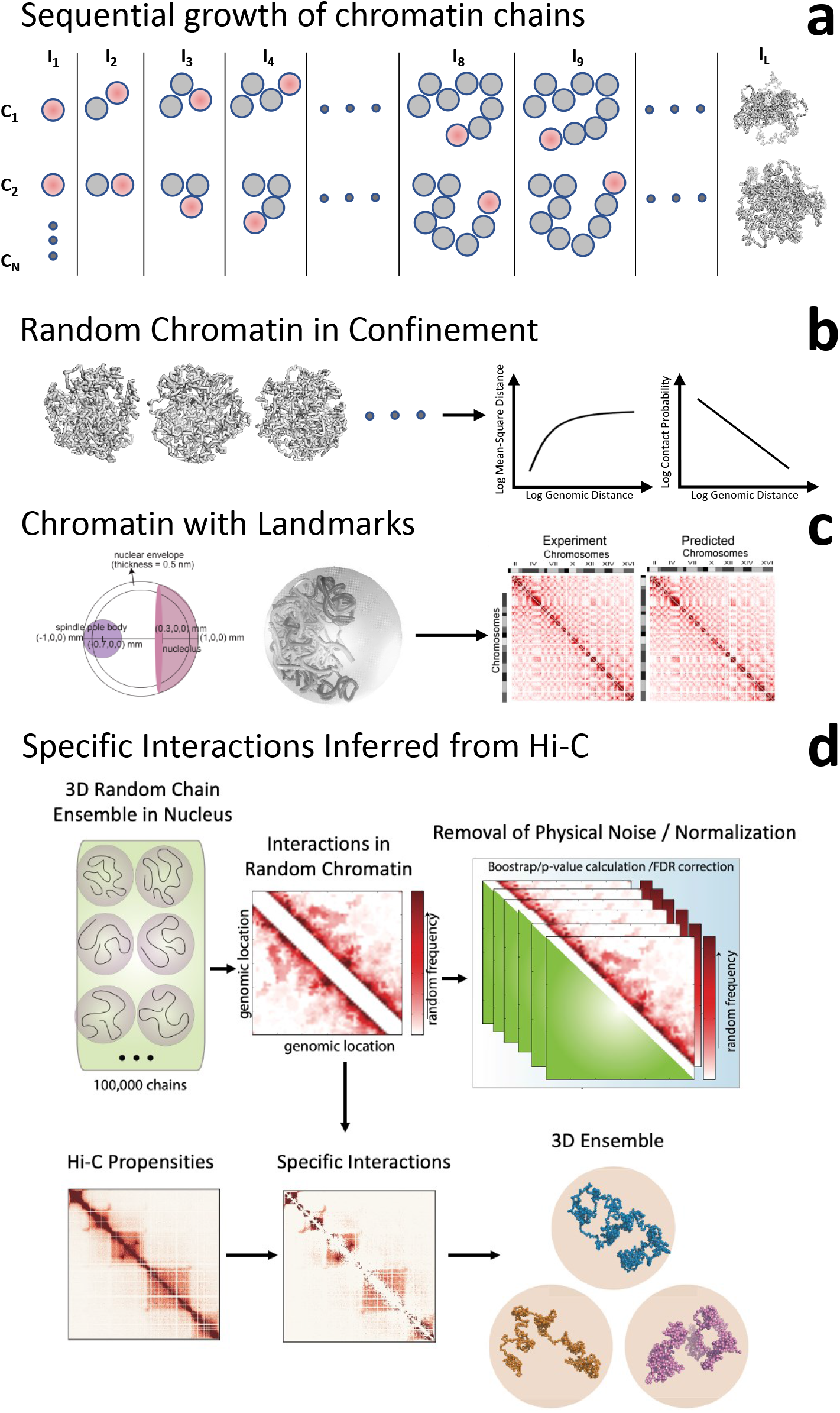
Minimalistic 3D self-avoiding polymer model. **(a)** Illustration of sequential polymer growth process. An ensemble of *N* polymer chains, *C*_1_… *C_N_*, is grown by iterative addition of self-avoiding monomer units until target length *L* is reached. **(b)** A confined ensemble of random polymers gives rise to the characteristic scaling relationships of mean-square distance and contact probability with genomic distance. **(c)** An ensemble of confined random polymers with nuclear landmarks (telomere and centromere attachments) imposed reproduces the heatmap of Hi-C measured frequencies, although no Hi-C information was used (from (38)); **(d)** The ensemble of confined random polymers can be used to remove spurious interactions in Hi-C measurements and to identify specific interactions, which have significantly elevated interaction frequencies. Upper subpanel illustrates that simulated random polymers, when aggregated, exhibit a heatmap of random interactions. The lower subpanel shows that spurious interactions can be removed from the experimental Hi-C maps through bootstrapping and FDR correction of the random heatmap, resulting in the identification of a small set of specific interactions. These specific interactions can then be used to fold chromatin into 3D structures (from (42)).

We next discuss recent findings using this minimalistic approach. First, their generic behavior without invoking Hi-C-derived information: experimentally measured chromatin scalings arise naturally, and incorporating telomere and centromere tethering results in 3D chromatin ensembles that reproduce Hi-C measured interactions in budding yeast.

### Nuclear Confinement Is Intrinsic to Chromatin Scaling Behavior

For two loci separated by a genomic distance *s*, their mean-squared distance *R*^2^(*s*) and contact probability *p*(*s*) scale characteristically. Specifically, *R*^2^ (*s*) ~ *s*^2*ν*^, with the exponent *ν* = 0.33 (46) and with *R*^2^(*s*) further plateauing at longer *s*. The contact probability scales as 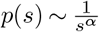, with the exponent *α* = 1.08 (1). Fractal globule was thought responsible for these behaviors (1), although subsequent studies suggested otherwise (47).

Imposing only nuclear volume confinement, randomly-generated ensembles of self-avoiding 3D chromatin polymers exhibit both scaling relationships and plateau behavior (Fig 2b) (38). These results show: 1) random and self-avoiding polymers in confinement give rise to scaling relationships and 2) nuclear size is a major determinant of chromatin folding. Indeed, nuclear morphology, such as a large ratio of nuclear to cytoplasmic volume, is likely important for the broad cellular reprogramming capabilities in stem cells (48). Further, nuclear size is often associated with diseases such as cancer (48).

### Nuclear Confinement With Landmarks Explains Yeast 3D Genome

In budding yeast, centromeres and telomeres are tethered to the spindle-pole body and nuclear envelope, respectively. Incorporating physical tethering of these nuclear landmarks into a basic polymer model was sufficient to give rise to the preferential localization of functional loci in the nucleus (49, 50). The ensemble of confined chromatin polymers with landmarks reproduced intra- and inter-chromosomal Hi-C interaction frequencies (Pearson *R* > 0.81 and *R* > 0.91, respectively) (43, 49, 50) (Fig 2c). In addition, centromere tethering was found to be responsible for inter-chromosomal interactions (43). Furthermore, fragile sites spatially cluster together (43). These findings demonstrate that ensembles of self-avoiding polymers in confinement, along with simple landmarks, can explain many Hi-C frequencies observed in yeast. There are significant implications for mammalian cells with analogous tetherings: attachment of heterochromatin to lamin, and association of actively transcribed regions to nuclear speckles (51–54). It would be interesting to quantitatively assess the extent to which these tetherings contribute to genomic 3D spatial organization, their functional consequences, and whether fragile sites similarly congregate spatially.

## A Small Set of Specific Interactions Can Drive Chromatin Folding

The results from the yeast polymer model require no Hi-C inputs. This suggests that many measured Hi-C interactions are due to generic effects of confined polymers: many Hi-C interactions occur due to random collisions of chromatin fibers confined in the cell nucleus (38, 42, 55). However, a small number of interactions occur at significantly elevated frequencies than would be expected in a randomly-generated ensemble of confined chromatin polymers. These over-represented interactions are known as *specific interactions* (42). In Drosophila, they constitute only 5 – 7% of measured Hi-C interactions. While their numbers are small, they appear to be critical in maintaining chromatin architecture, as simulations show that these interactions are sufficient to fold chromatin at both a locus and whole chromosome level. We discuss these below.

### A Small Fraction of Specific Interactions Captures Overall Hi-C Pattern

Specific Hi-C interactions emerge after filtering of spurious interactions which occur due to generic effects of polymer connectivity, self-avoidance, and nuclear confinement. To filter these spurious interactions, one compares the frequency of a Hi-C contact against the corresponding frequency in a simulated ensemble of randomly-generated polymer contacts (Fig 2d). Specific Hi-C contacts will have high frequencies meeting statistical significance and hence unlikely to have resulted from random collisions (38, 42). In Drosophila, analysis of 10 loci showed that only 5 – 7% of Hi-C interactions are specific (2.0 – 2.3 × 10^6^ out of 35 – 42 × 10^6^ Hi-C interactions); however, they capture the overall Hi-C pattern (Fig 3b) (45). Furthermore, there are clear changes in specific interactions during embryo development, such as reductions in active-inactive interactions and increases in inactive-inactive interactions (45). These changes, however, are undetectable without filtering of spurious Hi-C interactions (Fig 3b). This study illustrates that specific interactions reveal important temporal changes in chromatin structure not apparent in the measured Hi-C data.

**Fig. 3.**
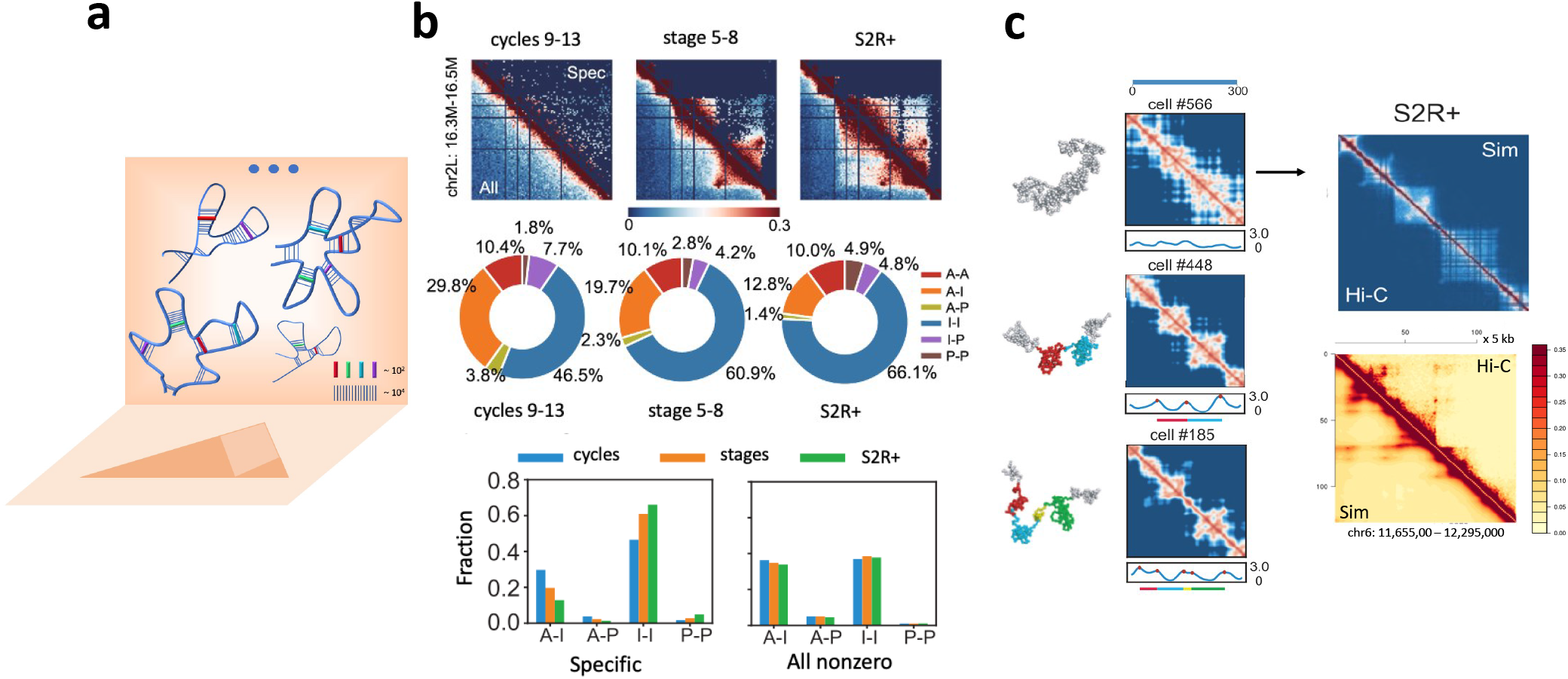
Chromatin can be folded using specific interactions. **(a)** Overview. A small set of interactions (thick colored sticks, 10 – 100 for a typical locus of 1 – 2 Mb) can fold chromatin in individual cells, where they appear in different combinations. Their aggregation gives rise to the 2D projection of the Hi-C heatmap. The Hi-C measured chromatin interactions (~ 10^4^) include both specific (colored sticks) and likely-biologically relevant, as well as bystander interactions (blue vertical lines); **(b)** Specific interactions in chr2L: 16.3̮16.5 Mb of Drosophila (upper triangles) capture the overall Hi-C interaction patterns (lower triangles) of cells at pre-MBT stages (cycles 9̮13), post-MBT (stages 5 – 8), and S2R+. Pie charts and bar charts show % of different types of specific interactions (A: active, I: inactive, P: polycomb-repressed) in the three cell types. As the embryo develops, A-I interactions (orange in pie chart) decreases, while I-I interactions (blue in pie chart) increase. In contrast, no such changes are detected when all Hi-C interactions are considered (bar chart, right) (from (45)); **(c)** Examples of single-cell chromatin conformations (chr2L: 11.0̮ 12.0 Mb, 2 kb resolution, Drosophila S2R+ cells) and their spatial-distance heatmaps. When aggregated, the simulated population Hi-C heatmap (upper triangle, top right column) resembles the measured Hi-C heatmap (lower triangle, Pearson correlation *R* = 0.95). (bottom right column) Experimental (top triangle) (2) and simulated (lower triangle) Hi-C heatmaps of human GM12878 cells (Chr6: 11.65 – 12.29 Mb, 5 kb) reconstructed by aggregation of single-cell conformations (lower triangle) folded using a small set of ≤35 specific interactions (*R* = 0.98) (from (44, 45)).

### Specific Interactions Can Fold Chromatin in Drosophila

An important question about 3D chromatin is whether there is a set of critical interactions that drive chromosomal folding (Fig 3a) (42, 56). Provided they exist, identifying them will likely reveal crucial determinants of 3D chromatin organization and delineate the important interactions and corresponding chromatin structures which may be necessary for genomic function.

Taking only the specific interactions for a locus (5 – 7% of all interactions) as distance restraints to be imposed on the self-avoiding 3D model, locus conformations generated using a deep-sampling algorithm reproduce Hi-C heatmaps with high accuracy (10 loci of length 0.2-2 Mb at different developmental stages at 2 kb resolution, Pearson correlation *R* = 0.91-0.98, Fig 3b) (45). In addition, details of loops and known long-range interactions are recovered (45). Furthermore, whole chromosome X of Drosophila can be folded at 5 kb resolution solely using specific interactions (45). Similar success was reported for the human *α*-globin locus, where independent ChIA-PET measurements were predicted from chromatin folded by 5C-derived specific interactions.

While 3D chromatin ensembles can be constructed by several methods, the success of the minimalistic self-avoiding polymer model is compelling, as only elementary physical considerations of fiber density, nuclear volume, and ligation distance are the model input. There are no adjustable parameters, no chromatin state assignments, and no *a priori* assumptions on loop anchors. Further, the small number of specific Hi-C interactions emerge naturally after spurious Hi-C interactions are removed using an ensemble of random conformations under the same volume confinement. These successes suggest that a small number of interactions, not necessarily indistinct chromatin states with unclear biological interpretation *per se*, may be sufficient to generate functional 3D chromatin ensembles.

### An Even Smaller Set of Specific Interactions Can Fold Chromatin Loci in Human Cells

The existence of driver interactions was further explored in a study examining multiple TAD-bounded regions with ≥ 2 super-enhancers in human GM12878 cells (44). Remarkably,only 14–35 specific interactions are sufficient to fold 39 loci (480 kb – 1.94 Mb) into Hi-C consistent ensembles at 5 kb resolution (*R* = 0.970 ± 0.003) (Fig 3c, bottom right). These are enriched with functional associations and active marks (44). They represent 0.024 – 1.3% (median 0.67%) of 2,414 – 62,785 measured Hi-C interactions, and 0.7 – 11% (median 5.7%) of the 301 – 2,112 specific interactions (44).

Interestingly, while removal of cohesin complexes and associated loops can abolish TADs (57), looping interactions alone (as defined by HiCCUPS (2)) are insufficient to drive chromatin folding for 10 Drosophila loci analyzed in (45). We expect similar results in the human loci studied in (44), as only 0.1% of the interactions in this smaller set are on domain boundaries or are loops identified by HiCCUPS. These results suggest that looping interactions occurring at TAD boundaries alone do not drive chromatin folding. Rather, a small set of specific interactions can fold chromatin into structural ensembles that naturally exhibit TADs.

While the absolute minimum set of driver interactions is unknown, it is intriguing whether this finding is general for other loci. Further analysis of these key interactions will be worthwhile to define the genomic elements involved and elucidate their functional roles.

## TADs Form as a Byproduct of 3D Chromatin Folding Driven by Specific Interactions

Topologically associating domains (TADs) (9, 10) are important structural units in our current understanding of 3D genome organization. However, their origin and role in genome function is of considerable debate (58–61). Minimalistic polymer models have shed some light on these.

### Proto-TADs Exist in Random, Volume-Confined Chromatin

When random chromatin polymers are confined within the nuclear volume, 3D domain structures with heightened intra-domain interactions arise naturally (Fig 4a)(38). These domains, called *proto-TADs*, are uniformly distributed along the genome (Fig 4c) (45), and appear frequently: on average, they cover ~21% of the random chromatin polymer (38). The entropic origin of TAD formation was also pointed out in (62). These results show that volume confinement induces a chromatin folding landscape with a propensity for TAD formation. This may greatly simplify the task of TAD formation during evolution: strategically placed protein factors may alter the folding landscape sufficiently so certain pre-existing proto-TADs are probabilistically favored and become fixed into TADs (Fig 4a).

**Fig. 4.**
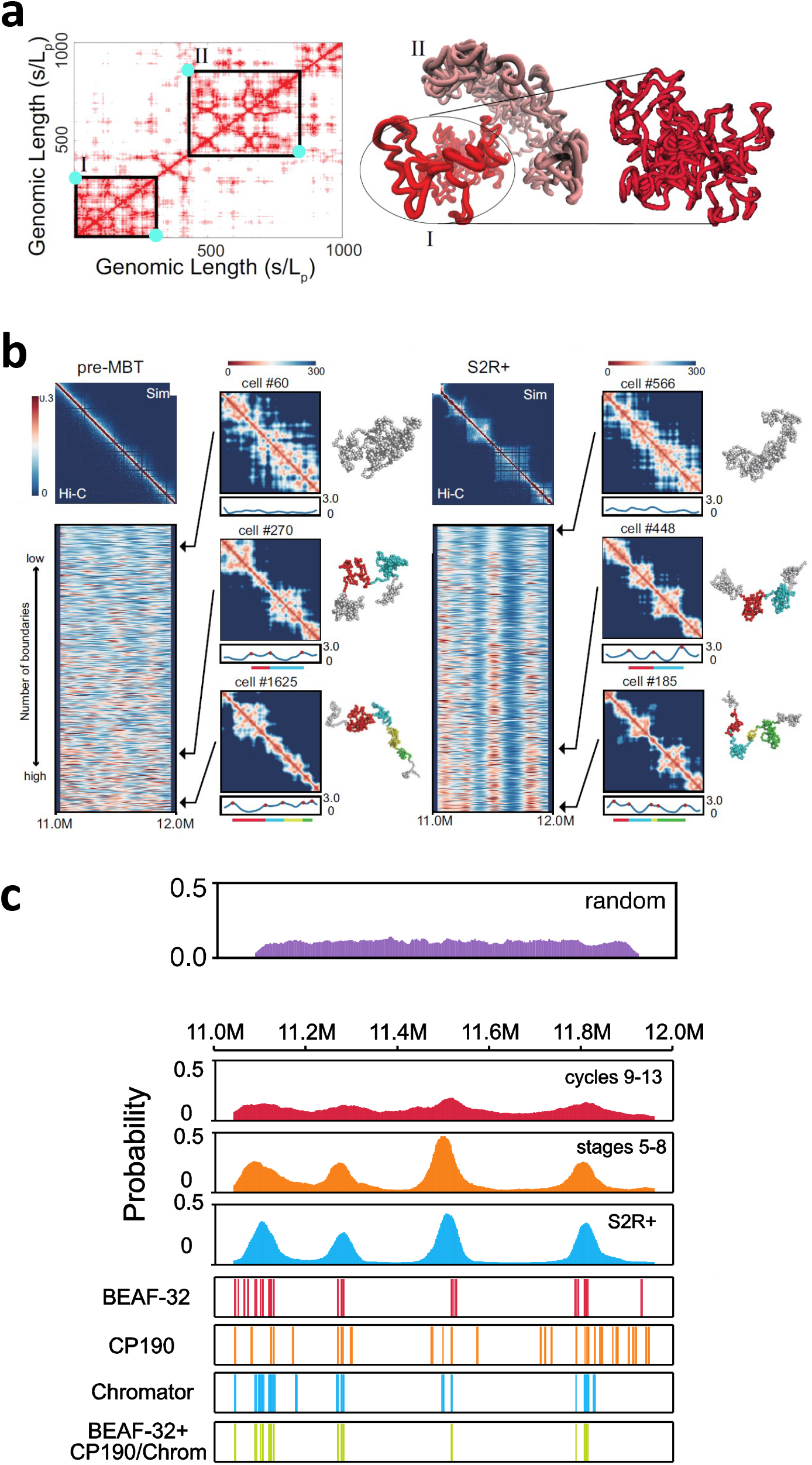
TAD formation as a byproduct of 3D chromatin folding driven by specific interactions. **(a)** Nuclear volume confinement induces a folding landscape where random polymers form domain-like 3D structures (right, domain I and II) which correspond to TAD-like patterns (left) in the 2D distance map (from (38)). Strategically placed protein factors (cyan dots) at the TAD boundaries for loop-formation may sufficiently tilt the chromatin folding landscape such that pre-existing proto-TADs are favored and become fixed; **(b)** Aggregated single-cell conformations of pre-MBT cells do not form population TADs, while those in S2R+ cells do. However, single-cell conformations of pre-MBT cells do form TAD-like structures, as also seen in single-cell spatial-distance heat maps. Boundary strength profiles of 5,000 conformations show most conformations in pre-MBT cells have ≥ 1 domain(s), but their boundaries have highly variable positions. In contrast, S2R+ cells have fixed boundaries; **(c)** Distributions of domain boundary probabilities along genomic position. Domains do exist in random polymers and are uniformly distributed with no preferred positions. Boundary probabilities increase at preferred positions as embryo cells develop. These boundary positions predicted from 3D single-cell conformations are preferentially localized at insulator binding sites such as BEAF-32, Chromator, and CP190.

### TAD Formation Is Driven by a Small Number of Specific Interactions

Studying TAD formation during embryo development has provided important insights. For a 1 Mb region of Drosophila, TADs are found in Hi-C maps of later cells (stages 5 – 8 and S2R+) but not in earlier pre-MBT cells (Fig 4b) (45). This pattern is also observed in modeled 3D conformations, which reproduce Hi-C maps at all stages (Fig 4b). Specifically, TAD-like structures are found in a large portion (60%) of individual S2R+ cells (Fig 4b), consistent with imaging studies (6, 63). However, despite there being no TADs in the Hi-C maps (Fig 4b, left), a large fraction (54%) of predicted single-cell chromatin conformations of early embryos contain TAD-like structures (Fig 4b). This paradox of TAD-like structures appearing in modeled individual cells but absent in population Hi-C can be understood from structural analysis of modeled single-cell chromatin conformations. Random polymers already possess TAD-like structures (Fig 4a) that are distributed uniformly (Fig 4c, top). At early embryo stages, simulations show TAD-like structures appear in individual cells, but with highly variable boundary positions and sizes (Fig 4b, left); reminiscent of random chromatin polymers and hence no TADs are detectable when aggregated. At later embryo stages, Hi-C domain boundaries become sharply peaked at distinct positions; correspondingly, aggregation of simulated single-cell conformations reproduces well-defined TADs with sharp boundaries (Fig 4c). These peaks are likely due to increased insulator binding: at TAD boundaries predicted from 3D conformations, binding of insulator complexes BEAF-32 and Chromator are highly enriched (Fig 4c) (45). Furthermore, boundaries in pre-MBT embryos associated with genes expressed zygotically before MBT (64) are in excellent agreement with boundary probabilities in modeled single-cell conformations (45).

These results suggest that TADs largely arise because of 3D chromatin folding driven by a small number of specific spatial interactions. The polymer folding landscape induced by the nuclear volume is already prone to form proto-TADs. With gradual introduction of strategically placed protein mediators of spatial interaction during development, TADs become favored, fixed, and appear as 2D patterns in population Hi-C. TADs are a byproduct of 3D interactions induced during folding, rather than a cause of 3D genome organization. This is consistent with recent findings that there is no simple relationship between TAD structures and gene expression (58–60).

## Minimalistic Single-Cell Chromatin Models Quantify Chromatin Heterogeneity and Uncover Many-Body Units

Hi-C reports only population averages, which hinders the detection of functional cellular subpopulations that may be important for developmental progression (45). Similarly challenging is detection of *many-body units*, three or more genomic regions simultaneously co-interacting within an individual cell, likely important for super-enhancer condensation (65, 66). Experimental single-cell techniques can address these issues but are limited in genomic coverage and sequencing depth (67). Alternatively, minimalistic chromatin models can now invert high-quality, population Hi-C into single-cell conformations whose aggregation is consistent with Hi-C measurements. We next briefly explain how this is accomplished (Fig 4a).

Minimalistic modeling of single-cell 3D chromatin must first predict which specific interactions identified from population Hi-C are present within an individual cell. Once specific contacts in individual cells are identified, 3D conformations are generated following the self-avoiding polymer growth approach (Fig 2a), with a distance restraint placed between loci of assigned specific contacts (Fig 5a). Central to this method is characterizing how certain interactions may cooperatively induce or exclude formation of other interactions and then account for these geometric dependencies when predicting cooccurrence of specific contacts within individual nuclei (44). This is accomplished through extensive polymer simulations using a Bayesian generative model (44). An alternative approach is described in (45). We next discuss the accuracy of these single-cell models as well as insights gained from them.

**Fig. 5.**
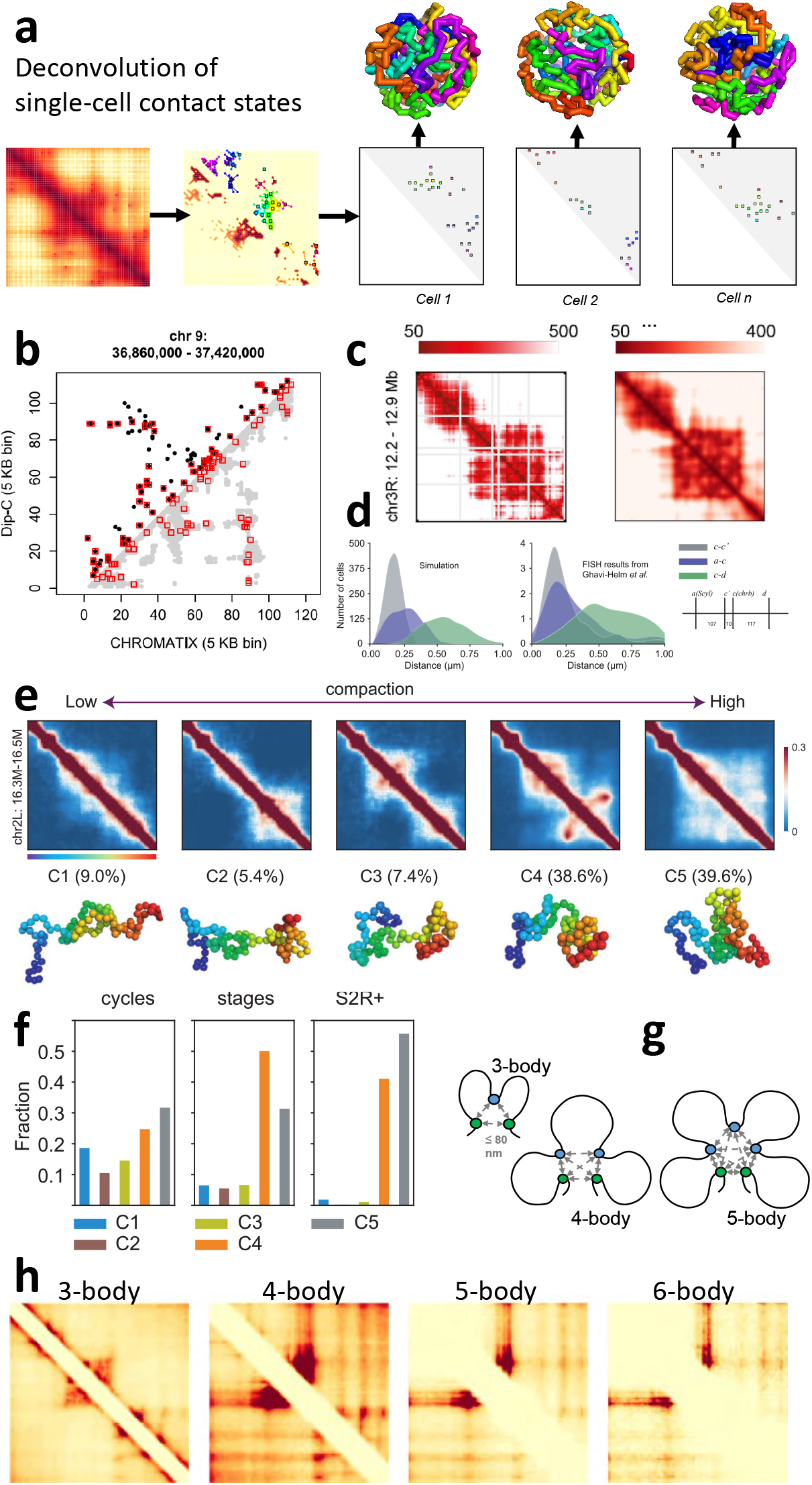
Single-cell chromatin conformations folded by specific driver interactions are accurate and reproduce Dip-C, imaging, and FISH studies. They can quantify chromatin heterogeneity and uncover interacting many-body units. **(a)** Illustration of inverting Hi-C measurements into an ensemble of single-cell chromatin conformations. Specific interactions are first identified from population Hi-C (see also Fig 2). Single-cell contact states (right) are then obtained through Bayesian deconvolution. Each contact state is then used to generate single-cell chromatin conformations, whose aggregation reproduces the poulation Hi-C heatmap. **(b)** Comparison with Dip-C single-cell data (GSE117874) (68). Pairwise contacts from a representative Dip-C single-cell locus (upper triangle, black dots) and the corresponding modeled single-cell locus conformation (lower triangle, gray dots). Contacts present in both models are outlined in red (from (44)). **(c)** Correspondence between distance maps of a modeled single-cell conformation (chr3R: 12.20̮12.90 Mb of Drosophila S2R+ at 2 kb resolution) with a conformation constructed from imaging studies in Mateo et al (63) (*R* = 0.75) (from (45)). **(d)** The distributions of spatial distances between chrb and Scyl genes and two other control regions in Drosophila derived from simulated single-cell chromatin conformations (Left) are highly consistent with DNA FISH measurements (Right) (59)(from (45))). **(e)** 3D single-cell chromatin conformations in a polycomb-repressed region of Drosophila can be grouped into 5 clusters, whose representative conformations are also shown. **(f)** The distributions of the 5 clusters in cell types of pre-MBT(cycles 9̮13), post-MBT (stages 5̮8) and S2R+. Collectively, these conformations reproduce the measured Hi-C heatmaps. **(g)** Diagrams of 3-, 4-, and 5-body chromatin interactions. All have pairwise Euclidean distance of ≤ 80 nm. The principal loop is the longest loop (in bp) among chromatin regions forming a many-body interaction. Green dots represent anchors of principal loops. **(h)** Principal loop heatmaps of *k*-body interaction units for the TAD (arrowhead) region containing the SH3KBP1 locus (chr X: 19,560,000̮20,170,000). Principal loop interaction frequencies are captured from deconvolved single-cells after aggregation (from (44, 45)).

### Single-Cell 3D Chromatin Is Accurately Modeled from Population Hi-C

Individual chromatin conformations generated are in excellent agreement with experimental singlecell measurements (44, 45). This concordance is seen when comparing with both single-cell imaging and Dip-C measurements (Fig 5b–5c). While reconciling FISH and Hi-C measurements is challenging (69), aggregation of individual 3D chromatin conformations can also reproduce the distance distributions of genomic regions by 3D-FISH (45) (Fig 4d).

Overall, minimalistic 3D polymer modeling can quantitatively invert statistical patterns in Hi-C heatmaps into highly-informative 3D single-cell chromatin conformations. The modeled conformations are not restricted in genome coverage, thus bridging the gap between population Hi-C and single-cell studies (6, 7, 63, 68, 70). This opens new avenues for modeling how genomic interactions in individual cells are related to cellular functions such as cis regulation of gene expression and replication.

### Single-Cell Chromatin Models Quantify Chromatin Heterogeneity

A cell population may have functional subpopulations with distinct chromatin conformations. However, it is not possible to directly inspect chromatin heterogeneity and identify subpopulations from Hi-C heatmaps. This requires an ensemble model of single-cell conformations with a properly defined distribution and thorough sampling.

Such single-cell 3D ensembles can now be generated (33, 42, 45). In Drosophila, chromatin heterogeneity of a polycomb-repressed region was quantified using 5.0 × 10^4^ single-cell conformations (45), which are found to form five clusters (Fig 5e). Their occupancies change dramatically as the embryo develops: evenly populated at an early stage, but two clusters dominate at the later S2R+ stage, accounting for *>*96% of the conformations (Fig 5f). Conformational subpopulations have also been quantified in single-cell models of the *α*-globin locus (42): K562 cells are homogeneous with a single cluster accounting for 97% of the cells, whereas GM12878 cells have many clusters and are far more heterogeneous.

The ability to characterize chromatin heterogeneity allows quantification of cellular subpopulations according to their intrinsic 3D chromatin structures. This will facilitate understanding of the relationship between genome structure and function. Important questions to be examined include: whether there are structural scaffolds facilitating *cis*-regulatory control of gene expression, are these scaffolds related to the distinct conformations representing cellular subpopulations, and can such a connection be substantiated with single-cell transcriptomics.

### Many-Body Units and Genome Function

Many-body (≥ 3) or multivalent spatial interactions likely play important roles in condensing super-enhancer (SE) regions into a transcriptional apparatus (65, 66). However, Hi-C (1) records only population-averaged, pairwise genomic interactions and therefore obscures which many-body interactions are present within individual cells. Single-cell Hi-C (70), Dip-C (68), Tri-C (71), MC-4C (72), GAM (73), and SPRITE (54) have great promise in uncovering multivalent chromatin interactions, but are currently limited in sequencing depth, genomic coverage, or inability to resolve direct versus indirect spatial interactions. Furthermore, it is challenging to evaluate if these spatial relationships are significant or simply explained by elementary physical effects of polymer connectivity, volume exclusion, and nuclear confinement.

The recent CHROMATIX method allows identification of specific many-body (≥ 3) interactions from Hi-C (44). Extending deep-sampling methods of (38, 42), it folds chromatin through fractal Monte Carlo sampling (44, 45, 74) and utilizes a Bayesian deconvolution approach to identify specific many-body units which are: i) fully spatially interacting, where all participating loci are within a Euclidean distance threshold (Fig 4g), and ii) not arising from aforementioned elementary polymer effects.

The functional landscape of many-body interactions of 39 transcriptionally-active TADs in human GM12878 cells (2) was constructed using CHROMATIX. Many-body units were found to occur frequently in these euchromatic loci. Compared to randomly formed many-bodies, specific many-bodies were enriched in promoters, enhancers, and superenhancers. In addition, anchor loci of principal loops - the longest spanning loops within many-body units (Fig 4g), exhibit banding patterns when projected as a 2D-heatmap and are enriched in super-enhancers. These results show principal loops likely bridge enhancers and promoters to enable spatial coupling of functional regions. As reported in a recent Drosophila study (75), there is now emerging evidence that analogous multi-way interactions among enhancers and promoters are pre-formed in early embryo development and then become activated or repressed during developmental progression.

Principal loop anchors of specific many-bodies can be directly predicted using 1D biomarkers (44). DNase accessibility was found to be the most predictive biomarker. POLR2A occupancy and nuclear fraction RNA abundance are also important predictors, indicating these specific many-bodies may help facilitate transcription. This is consistent with a subsequent study (76) proposing RNA accumulation as a mechanism of microphase separation, resulting in co-segregation of transcriptionally-active chromatin. CTCF and cohesin subunit RAD21 were modestly predictive of specific principal loops, indicating that while loop extrusion (47, 77) may occur within the examined TAD regions, there are likely other important mechanisms at play in the formation of many-body units enriched in functional elements.

## Outlook

Minimalistic self-avoiding 3D chromatin modeling with few tunable parameters has revealed insights into 3D genome organization: 1) chromatin scaling rules are a result of volume-confined polymers, 2) chromatin folding at many loci is driven by a small number of specific interactions, 3) TADs emerge from ensembles of single-cell chromatin folded according to these small number of specific interactions, 4) heterogeneous structural scaffolds help define intrinsic cellular subpopulations whose relative representations are likely important to developmental progression, and 5) the extent and functional roles of many-body spatial interactions in enhancer-rich regions, which are enabled by principal loops. These findings point to several interesting future directions. A first task is to identify specific driver interactions in a wide range of loci. This can be done broadly for different cell types to define the genomic elements involved and their functional roles. Furthermore, it will be highly informative to assess which specific interactions are conserved and which are unique across different cell types, so the structure-function relationship of chromatin can be characterized at the tissue level.

In addition, there may be better representations of 3D chromatin structure. While the relationship of function and form is of extraordinary importance to understanding 3D genome organization, the prevailing representations of 3D genome structure in terms of TADs and subTADs (78, 79) are inadequate, as they are indirect 2D projections of 3D chromatin and more direct representations may be feasible. These representations could be based on the set of interactions that drive chromatin folding, the principal structures within clusters of 3D conformations, or by characterization of the folding mechanisms among pioneering interactions and other functional interactions enabled by them. Such new representations can stimulate investigation into the most relevant molecular players, their interactions, and the biological processes involved.

A practical utility of such a representation is to allow targeted perturbation of 3D genome organization. For genomic regions that are poorly characterized, current perturbations may involve reversal/deletion of large (≥ 100 kb) genomic intervals such that many elements are disrupted, and their accumulative effects detected. If a small set of critical driver interactions can be identified for a specific locus, it would facilitate experimental perturbations of much smaller intervals to assess the extent of 3D chromatin alteration, and thereby precisely identify regions important for nuclear organization, enabling further investigation of their molecular functions. Such a perturbation strategy may be generally applicable to any arbitrary locus. The small number of specific interactions identified in (42, 45, 74) suggest this may be the case. Furthermore, functional characterization of perturbed regions may allow *a priori* prediction of resulting phenotypic changes which may then be experimentally validated; this is similar to the process used in discovery of *cis* elements controlling DNA replication (80).

Additionally, it should be possible to define the functional landscapes of many-body spatial chromatin interactions in different cells and tissues. With the identification of the participating functional elements and their principal loops, we will gain better understanding on how novel loop anchors can spatially coalesce multiple functional elements to form a coherent apparatus for transcription. A global understanding of many-body interactions is indeed feasible, as demonstrated by the large-scale study of a set of enhancer-rich loci (44).

Lastly, we have a new means to investigate how chromatin structure can provide the physical basis for cellular activity. We can asssess how cellular programs as defined by singlecell transcriptomes correspond to single-cell 3D chromatin conformation. The ability to quantify structural heterogeneity may allow us to delineate functional cellular subpopulations based on their shared chromatin folds. For example, we can assess correspondence between different structural clusters of chromatin and different types of cellular behavior, and whether certain 3D scaffolds are required for specific cellular states. The analysis of time-evolving patterns of chromatin clusters may further shed light on embryonic development as seen in Drosophila (Fig 4e–4f) (45).

## Conclusions

With powerful polymer models that can transform 2D Hi-C interaction maps into ensembles of single-cell 3D chromatin conformations, we expect to gain further understanding of the genome structure-function relationship. Emerging frontiers include: 1) relating ensembles of spatial chromatin structures to cellular phenotypes such as gene expression and gene usage; 2) establishing the structural basis and identifying spatial motifs for different cellular states that are consistent with experimentally measured single-cell transcriptomics; and 3) relating transcriptional heterogeneity to chromatin structural heterogeneity to improve understanding of embryogenesis and cellular reprogramming (81). We expect that integrated modeling and experimental studies will play important roles in investigating these important questions.

## Acknowledgements

We thank Drs. Kostas Chronis, Dan Czajkowsky, Gamze Gursoy, Amy Kenter, Ao Ma, Zhifeng Shao, Jan-Hendrik Spille, Qiu Sun, and Wei Yang for their critical reading of this manuscript. We thank Hammad Farooq and Lin Du for their assistance in preparation of this manuscript. This work is supported by NIH Grant R35 GM127084.

## Summary

- A minimalistic, self-avoiding 3D chromatin model with no adjustable parameters can transform population Hi-C into high-resolution, single-cell chromatin conformations;
- Chromatin folding at many loci is driven by a small number of specific interactions;
- TADs form as a byproduct of 3D chromatin folding driven by specific interactions;
- Cell subpopulations equipped with different chromatin structural scaffolds are developmental stage-dependent;
- Characterization of the functional landscape and epigenetic marks of many-body units which are simultaneously spatially co-interacting within enhancer-rich, euchromatic regions.

